# RsmW is an iron-responsive 3’ UTR derived small regulatory RNA that contributes to biofilm formation and virulence of *Pseudomonas aeruginosa*

**DOI:** 10.64898/2026.09.03.749296

**Authors:** Rhishita Chourashi, Amanda G. Oglesby

## Abstract

*Pseudomonas aeruginosa* is an opportunistic Gram-negative pathogen that causes acute and chronic opportunistic infections. Progression to chronic infection requires upregulation of the Rsm small regulatory RNAs (sRNAs), which in turn induce the expression of virulence factors to promote chronic virulence phenotypes, including biofilm formation. Iron is a critical nutrient for survival and virulence of *Pa*, and host-mediated iron limitation leads to the induction of key virulence factors driving infection. Prior work showed that the RsmY and RsmZ sRNAs are modestly upregulated due to iron limitation in static growth conditions via PqsR. Here, we show that another Rsm sRNA, RsmW, is strongly induced by iron limitation, likely in a Fur-dependent manner. We identified an RNA processing event, which appears to be mediated by the RNAse E, resulting in cleavage of RsmW from the 3’ UTR of *pa4570* mRNA. We also report that RsmW is more stable than the full length *pa4570-rsmW* transcript as well as the processed *pa4570* mRNA. Surprisingly, we found that RetS, which down-regulates RsmY and RsmZ expression, enhances expression of RsmW under iron limiting conditions. We further demonstrate that RsmW plays a key role in biofilm formation under iron limiting conditions and is required for full pathogenesis in a *Galleria mellonella* infection model. Lastly, we observed similar iron-dependent regulation of RsmW in chronic infection isolates from cystic fibrosis (CF) sputum, underlining the potential importance of this regulation in the CF lung. Taken together, the results outline a new paradigm for sRNA-dependent iron regulation of *P. aeruginosa* pathogenesis.

**AUTHOR SUMMARY:** *P. aeruginosa* requires iron for its survival and virulence. Upon infection the mammalian host limits iron, which *P. aeruginosa* overcomes via numerous regulatory responses that play a central role in pathogenesis. He we report that iron starvation induces expression of the RsmW small regulatory sRNA, which regulates key virulence phenotypes that promote the progression of chronic infections in the CF lung. We show that RsmW is processed from the 3’ untranslated region (UTR) of the *pa4570* transcript, which is also induced upon iron starvation. Our studies also revealed regulatory events that are distinct from the previously characterized Rsm sRNAs, suggesting RsmW functions under distinct environmental conditions. Furthermore, we found that RsmW contributes to biofilm formation in flow cells, promotes killing in *Galleria mellonella* larvae, and is induced by CF isolates upon iron starvation. Taken together, our work demonstrates that RsmW is a novel iron-dependent regulator of chronic virulence traits in *P. aeruginosa*.

## INTRODUCTION

*Pseudomonas aeruginosa* is an opportunistic Gram-negative pathogen that causes acute and chronic lung infections in compromised populations, including individuals with cystic fibrosis (CF). *P. aeruginosa* also infects wounds in burn victims, diabetics, and surgical patients (1–5). In the CF lung, *P. aeruginosa* colonization initiates as an acute infection and gradually shifts to chronic, multi-drug resistant infection over years to decades (6). *P. aeruginosa* requires iron for growth and infection, yet the host innate immune system sequesters iron to limit microbial growth through a process termed nutritional immunity (7). *P. aeruginosa* overcomes iron limitation via iron starvation responses that allow it to scavenge host iron (7–10). These responses include the upregulation of genes for production and uptake of siderophores, which exhibit exquisitely high affinities for iron. Expression of iron uptake genes is regulated by the ferric uptake regulator (Fur), a global transcriptional repressor that is dependent on iron for DNA binding (11). Fur recognizes a consensus sequence, known as the Fur box, in the promoter region of iron responsive genes, thereby inhibiting RNA polymerase from transcribing iron uptake genes when cytosolic iron is replete. Fur regulation therefore balances *P. aeruginosa’s* requirement for iron with the toxic potential of this metallo-nutrient.

Many bacterial species also produce Fur-regulated small regulatory RNAs (sRNAs) that are expressed upon iron starvation (12). In *P. aeruginosa*, iron starvation leads to the induction of the Fur-repressed PrrF sRNAs, which pair with and prevent translation of mRNAs for non-essential iron-dependent proteins to mediate an iron sparing response (13–16). Our prior studies have shown that the PrrF sRNAs are required for *P. aeruginosa* iron homeostasis, twitching motility, and virulence (14,16). More recently, we showed that PrrF contributes to biofilm formation in iron-limiting conditions by promoting the production of alkyl-quinolone (AQ) metabolites produced through the Pseudomonas quinolone signaling pathway (17). Moreover, PrrF is expressed in clinical isolates from CF patients, supporting their role CF lung infection (18). Thus, the PrrF sRNAs are central mediators of *P. aeruginosa* iron homeostasis and virulence.

We previously showed that two additional sRNAs, RsmY and RsmZ, are modestly upregulated by *P. aeruginosa* upon iron starvation (19). Curiously, this regulation was observed when *P. aeruginosa* was grown under static but not shaking conditions, the former of which we posit is more representative of the *in vivo* environment. The RsmY and RsmZ sRNAs belong to the Rsm regulatory system that consists of two additional sRNAs (RsmW and RsmV) and the RNA binding proteins RsmA, RsmN, and RsmF (20–25). The Rsm sRNAs sequester the Rsm RNA binding proteins, which bind to GGA motifs in the Shine Dalgarno (SD) sequence of targeted mRNAs to prevent their translation (20,21,23,25). RsmY and RsmZ are upregulated by the GacS/GacA two-component system, which is mediates the switch from acute to chronic infection during CF lung infections (23,26). The Gac/Rsm pathway is repressed during acute infections by the RetS histidine kinase, the gene for which undergoes mutational inactivations that accumulate in many CF isolates of *P. aeruginosa* (27). While the contribution of RetS/GacS/Rsm pathway in the regulatory switch from acute to chronic *P. aeruginosa* infection is clearly established, the role of iron in this switch remains unclear.

The current study sought to determine the effects of iron on the expression of the RsmW sRNA. RsmW is transcribed as the 3’ UTR of the *pa4570* mRNA, and a consensus Fur box is located upstream of this locus. Accordingly, we show that iron starvation results in robust upregulation of RsmW and *pa4570*. We also show that the *rsmW-pa4570* transcript is cleaved into a stable RsmW sRNA and relatively less stable *pa4570* mRNA. Moreover, we provide evidence that RNase E contributes to this processing event. We further report that deletion of *retS* leads to reduced RsmW expression, an effect that is opposite of what is observed here and elsewhere for RsmY and RsmZ (28,29). Expression of RsmW is also strongly induced by iron starvation in clonal isolates from the lungs of individuals with CF, supporting its potential role in iron dependent regulation of *P. aeruginosa* pathogenesis. Lastly, we showed Δ*rsmW* is defective in flow-cell biofilm formation under iron starvation conditions and is attenuated in a *Galleria mellonella* infection model. Altogether, our findings indicate RsmW is regulated by distinct environmental signals from RsmY and RsmZ and outline a key role for this sRNA in *P. aeruginosa* biofilm formation and virulence.

## RESULTS

### Iron starvation induces expression of *pa4570* and RsmW

Sequence analysis revealed that the *pa4570_rsmW* locus is preceded by a Fur box, which is conserved at the *pa4570* locus in other pseudomonads (Fig. S1A). In contrast, the *rsmW* sequence is specific to *P. aeruginosa* (Fig. S1A.iii). We therefore investigated whether iron regulates *pa4570_rsmW* in *P. aeruginosa* by growing *P. aeruginosa* strain PAO1 for 18 h in DTSB medium with (high iron) or without (low iron) 100 µM FeCl_3_. We designed primers against the RsmW sRNA, the *pa4570 mRNA*, and the predicted *pa4570-rsmW* co-transcript for qPCR analysis (Fig. 1A). Importantly, the RsmW and *pa4570* primer sets also detect the *pa4570_rsmW* co-transcript (Fig. 1A). qPCR analysis revealed robust expression of RsmW and *pa4570* RNA levels in low iron conditions and full repression upon iron supplementation when cDNA was amplified by all three primer sets (Fig. 1A). Northern blot analysis using a 5’ biotin-labelled RsmW specific probe further demonstrated iron-repressed expression of RsmW (Fig. 1B, Fig. S1B). The RsmW-specific probe also detected the *pa4570_rsmW* co-transcript in the low iron condition (Fig. 1B). Together, our data indicate *pa4570_rsmW* transcript is strongly regulated by iron, likely in a Fur-dependent manner.

**Figure 1.**
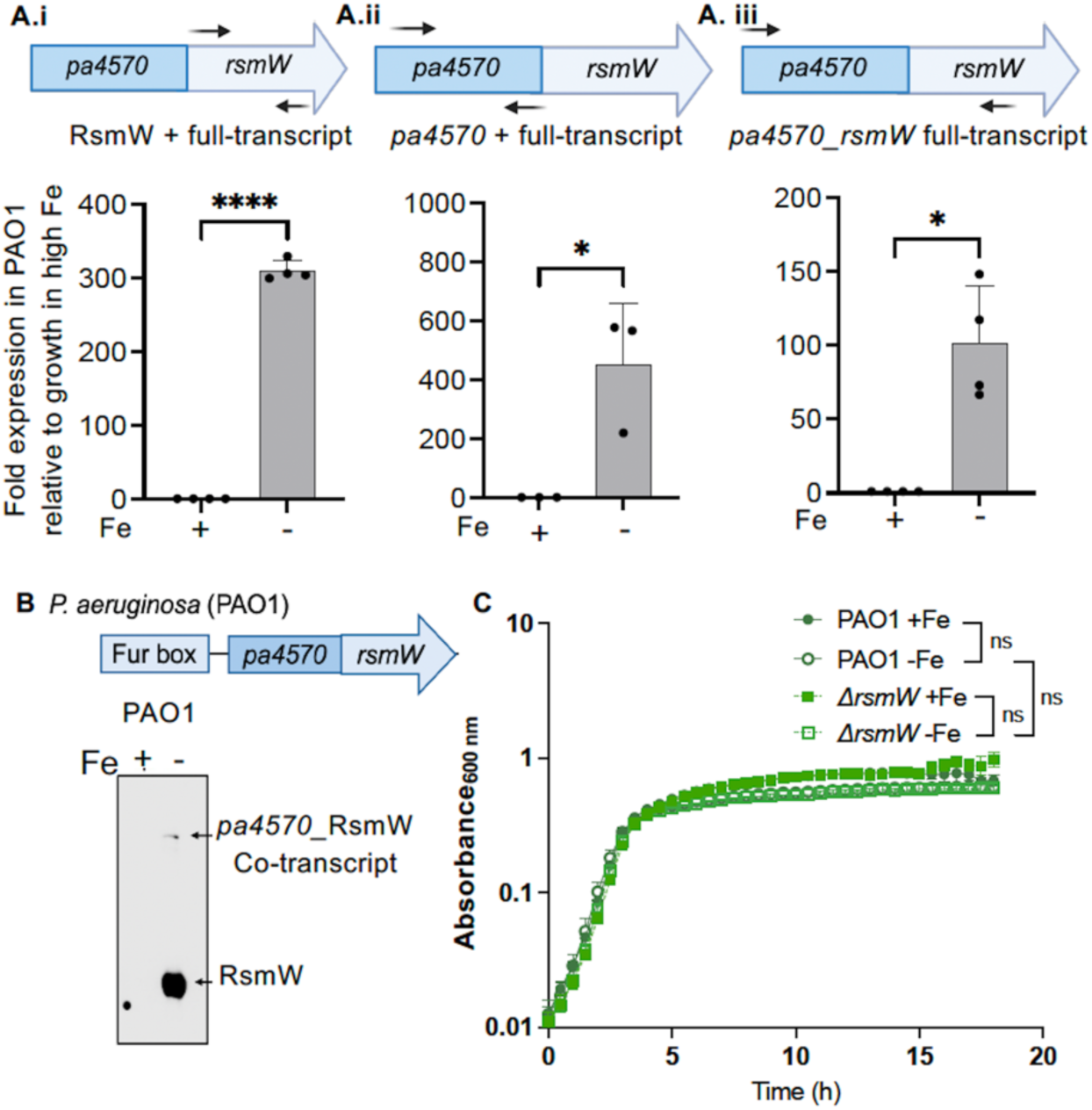
RsmW is induced under iron limiting conditions but is not required for growth. A) qPCR of i. RsmW, ii. *pa4570* and iii. *pa4570_rsmW*, using the primers indicated above each graph, was performed on cDNA from PAO1 cultures grown in high and low iron conditions. qPCR was performed with at least 3 biological replicates and 3 technical replicates per biological replicate. Each biological replicate is represented a black circle data point. B) Northern blot assays were performed using biotinylated oligonucleotide probes that detect RsmW. Northern blot image is representative of 3 biological replicates. C) The indicated strains were grown in 96 well plate for 18 h in DTSB media supplemented with or without iron. Results are represented as mean ± SD. Asterisks indicate a statistical difference as determined by T-test (A) or two-way ANOVA followed by Bonferroni’s multiple comparison test (C): * P < 0.01, **** P < 0.0001.

Since RsmW is induced by iron starvation, we questioned whether RsmW was required for optimal growth in low iron conditions. Accordingly, we analyzed growth of PAO1 and the Δ*rsmW* mutant in DTSB media with or without iron supplementation for 18 h. Both strains grew similarly regardless of iron supplementation conditions (Fig. 1C), indicating that loss RsmW does not impact growth during iron starvation.

### The RsmW sRNA is more stable than the *pa4570* mRNA

Prior work predicted that the RsmW sRNA is processed from the 3’ UTR of *pa4570* (20). To examine the stability of the co-transcript and cleavage products, PAO1 was grown in DTSB without iron supplementation for 18 h, at which point cultures were supplemented with 100 µM of FeCl_3_ to stop transcription, and RNA was harvested at various time points for northern blot analysis using probes against RsmW and *pa4570* (Fig. 2A). Our results indicate that RsmW is stable up to 40 min post iron supplementation, after which RsmW abundance showed a slight decline (Fig. 2B and Fig. S2, top panels). In contrast, the *pa4570_*RsmW co-transcript exhibited a rapid decline; this decline was more apparent with the RsmW probe than with the *pa4570* probe, likely due to the more abundant RsmW sRNA titrating away the probe (Fig. 2B, compare top and bottom panels). Notably, the transcripts detected by the *pa4570*-specific probe rapidly disappeared and laddering was observed (Fig. 2B and Fig. S2, bottom panels). These data indicate that the processed RsmW sRNA is more stable than the processed *pa4570* mRNA.

**Figure 2.**
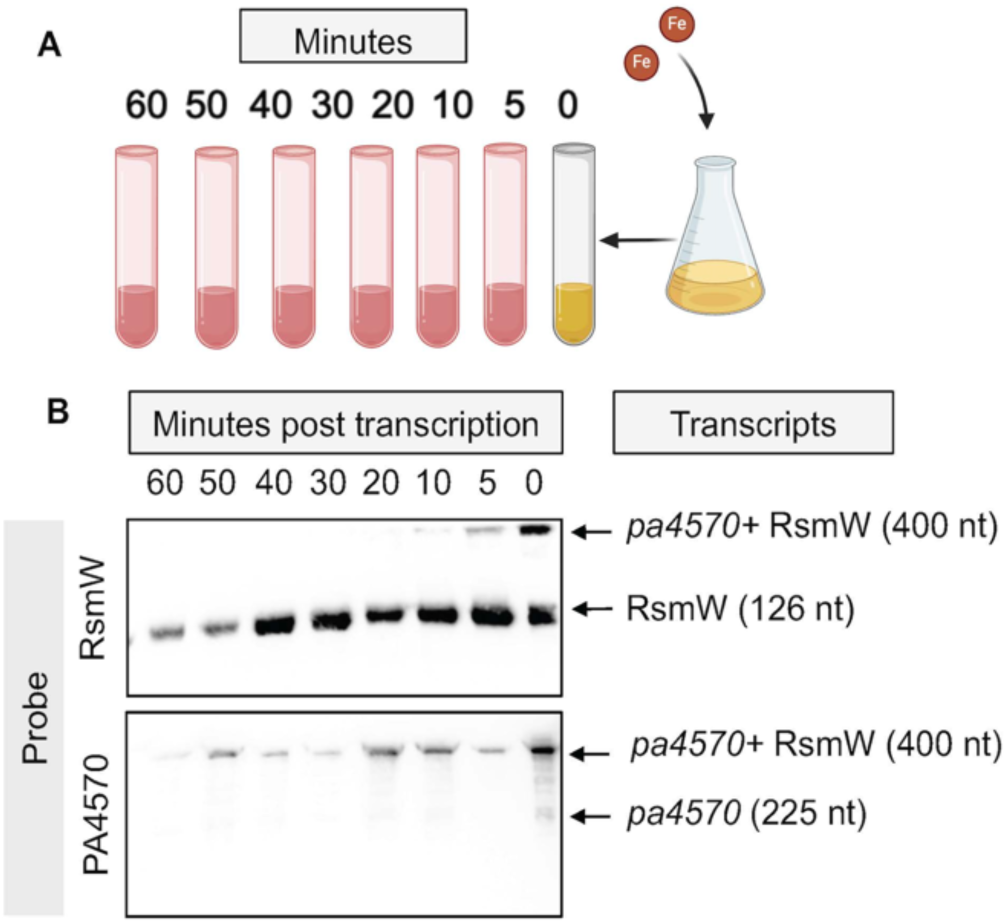
RsmW is robustly stable post-transcription and cleavage. A) Schematic of method used for stability assays. B) PAO1 cultures were grown in DTSB for 18 h without Fe and cultures were harvested at (0, 5, 10, 20, 30, 40, 50, 60) min after iron (Fe) supplementation as mentioned in the panel A. RNA was isolated and northern blot assay was performed using biotinylated oligonucleotide probes against either RsmW or *pa4570*. Northern blot images are representative of 3 biological replicates.

### RNAse E lacking its degradosome scaffold domain results in decreased processing of the *pa4570_rsmW* transcript

Towards understanding how the *pa4570_rsmW* co-transcript is processed to produce the *pa4570* mRNA and Rsm sRNA, we identified a putative RNAse E cleavage site within the *pa4570_rsmW* co-transcript (Fig. S3A). To this end, we quantified levels of the RsmW sRNA, *pa4570* mRNA, and *pa4570_rsmW* co-transcript in wild type PAO1 and an isogenic mutant that lacks 50bp of *rne* corresponding to the C-terminal region of RNase E (Fig. S3B), which serves as the scaffold for assembly of the RNA degradosome complex (30). PAO1 and Δ*rne 50bp c-term* were grown in DTSB with or without 100 µM of FeCl_3_ supplementation for 18 h, at which time RNA was isolated for qPCR analysis. Iron repressed expression of the *pa4570* and RsmW RNAs regardless of the primer set that was used. However, we observed a significant increase in the expression of the *pa4570_rsmW* co-transcript levels in Δ*rne 50bp c-term* compared to PAO1 under low Fe conditions (Fig. 3A), and a significant decrease of the RsmW sRNA in the Δ*rne 50bp c-term* under low Fe conditions (Fig. 3B). Similar to the co-transcript levels, analysis of the samples with the qPCR primers and probes specific to *pa4570* showed increased expression in Δ*rne 50bp c-term* as compared to wild type PAO1 (Fig. 3C), which may be due to increased stability of the co-transcript as compared to the processed *pa4570* mRNA. In turn, the decrease in RNA detected by the RsmW probe, which binds to both the processed RsmW sRNA and the *pa4570_rsmW* co-transcript, suggests that the RsmW sRNA is more stable than the co-transcript. Taken together, our data indicate that the RNAse E degradosome scaffold contributes to the RNA processing of *pa4570_rsmW* co-transcript.

**Figure 3.**
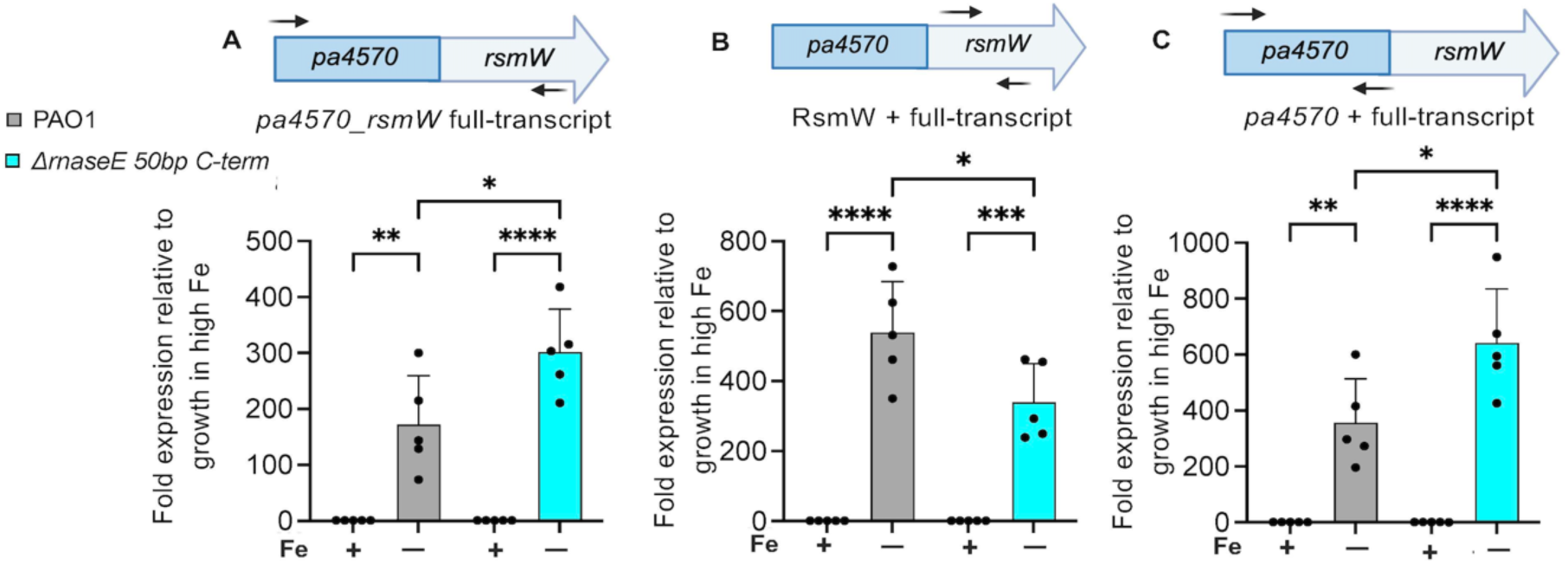
The RNase E–associated degradosome scaffold contributes to the processing of *pa4570_rsmW* co-transcript. The indicated strains were grown in DTSB media with or without iron supplementation for 18 h. Cultures were harvested, and RNA was isolated. cDNA synthesis was done, and qPCR was performed using primers specific for A) RsmW, B) *pa4570* and C) *pa4570_rsmW* co-transcript. 5 biological replicates and 3 technical replicates per biological replicate were analyzed. Each biological replicate is represented a black circle data point. Results are represented as mean ± SD. Asterisks indicate a statistical difference as determined by two-way ANOVA followed by Bonferroni’s multiple comparison test: * P < 0.05, ** P < 0.005, **** P < 0.0001.

### Deletion of *pa4570* results in reduced RsmW sRNA levels

Since the *pa4570_rsmW* co-transcript is processed into distinct RNA molecules, we determined whether loss of either of the genes would affect expression levels of the other gene. To this end, we generated Δ*pa4570* and Δ*rsmW* single mutant strains in the PAO1 background. Strains were grown for 18 h in DTSB supplemented with or without 100 µM of FeCl_3_ and analyzed by qPCR. These data showed a reduction in RsmW sRNA levels in Δ*pa4570* compared to PAO1 under iron starvation conditions (Fig. 4A), while loss of RsmW resulted in an insignificant reduction in *pa4570* mRNA levels (Fig. 4B). Thus, our data indicate that *pa4570* affects RsmW levels, though the origin of this effect is unclear, whereas *rsmW* has limited impact on *pa4570* transcript levels.

**Figure 4.**
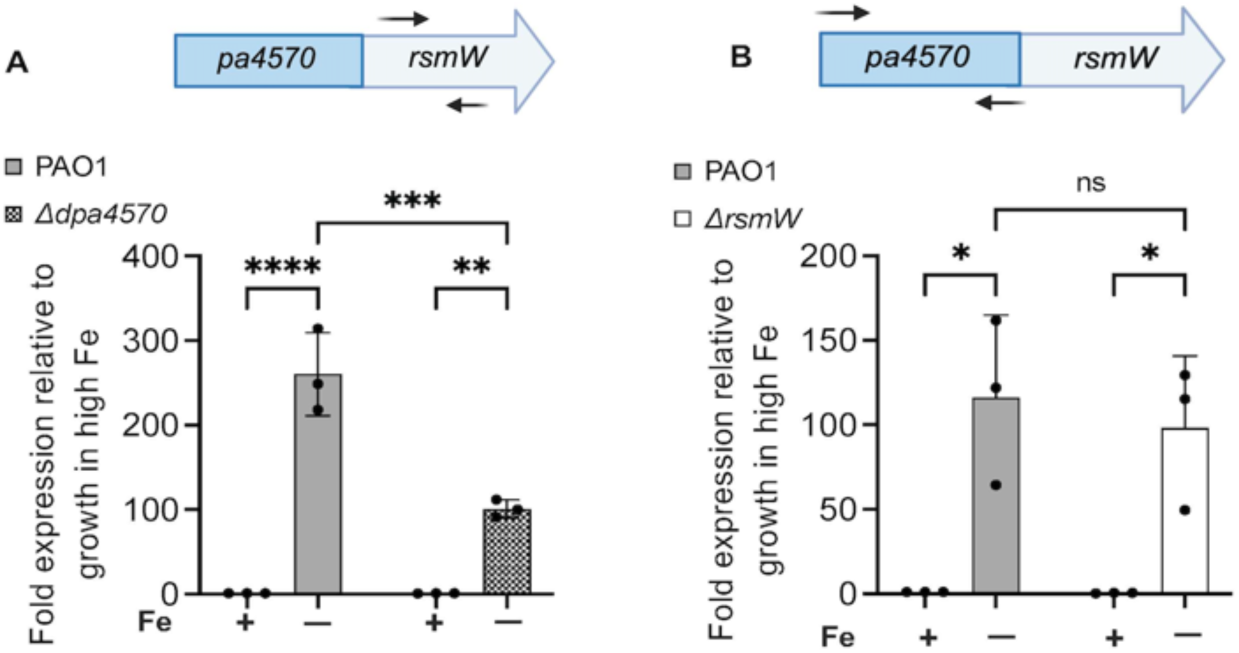
The *pa4570* gene sequence is required for full RsmW expression. The indicated strains were grown in DTSB media with or without iron supplementation for 18 h. Cultures were harvested, and RNA was isolated for cDNA synthesis and qPCR analysis using the RsmW (A) or *pa4570* (B) primers. 3 biological replicates and 3 technical replicates per biological replicate were analyzed. Each biological replicate is represented a black circle data point. Error bars indicate standard deviation (SD). Asterisks indicate a statistical difference as determined by two-way ANOVA followed by Bonferroni’s multiple comparison test: * P < 0.05, ** P < 0.01, *** P < 0.0005, **** P < 0.0001.

### RetS positively affects levels of the RsmW sRNA but not the *pa4570* mRNA

Since the two-component sensor kinase RetS is known to repress expression of the RsmY and RsmZ sRNAs via Gac/GacA two component system, we next determined the impact of RetS on RsmW expression. Accordingly, we used qPCR to compare expression of RsmW and *pa4570* in PAO1 and an isogenic Δ*retS* mutant grown in DTSB with or without iron supplementation. Notably, *retS* deletion resulted in decreased expression of RsmW (Fig. 5A), while having no effect on *pa4570* mRNA levels (Fig. 5B). This finding was surprising as both RsmW and *pa4570* are predicted to share the same promoter, suggesting the impact of RetS signaling on RsmW levels occurs post-transcriptionally. Moreover, the observation that Δ*retS* shows decreased RsmW expression is opposite of that observed here (Fig. 5C-D) and in prior work (26,31) for the RsmY and RsmZ sRNAs. These results further demonstrate that RsmW is subject to distinct signaling pathways than RsmY and RsmZ.

**Figure 5.**
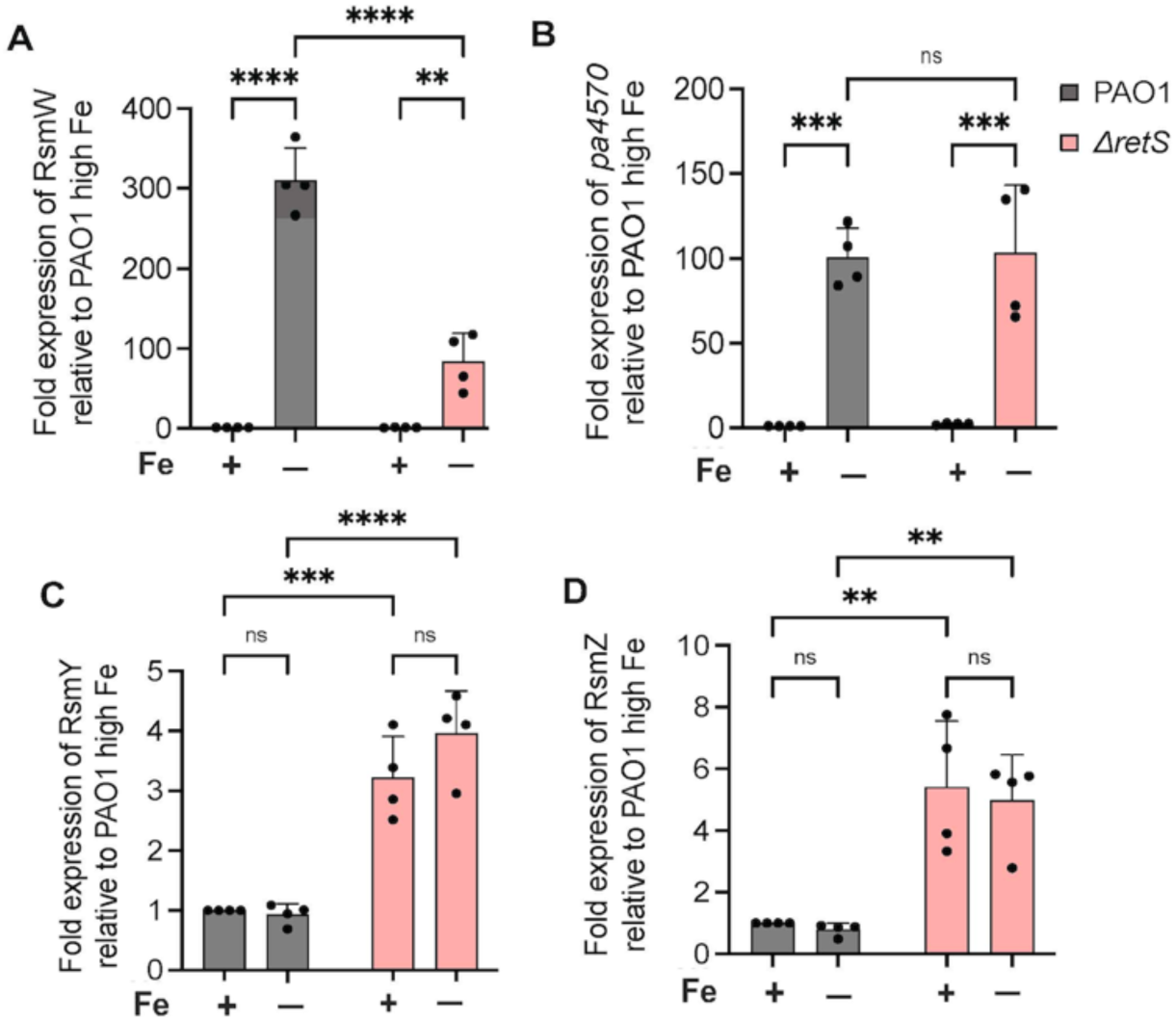
RetS differentially impacts levels of RsmW, *pa4570,* RsmY, and RsmZ. The indicated strains were grown in DTSB media with or without iron supplementation for 18 h. Cultures were harvested, and RNA was isolated for cDNA synthesis and qPCR analysis using primers specific for RsmW (A), *pa4570* (B), RsmY (C), and RsmZ (D) upon iron starvation between PAO1 and Δ*retS*. 3 biological replicates and 3 technical replicates. Each biological replicate is represented a black circle data point. Results are represented as mean ± SD. Asterisks indicate a statistical difference as determined by two-way ANOVA followed by Bonferroni’s multiple comparison test: * P < 0.05, ** P < 0.01, *** P < 0.0005, **** P < 0.0001.

### The RsmW sRNA is required for biofilm formation in low iron conditions

Prior work showed that RsmW expression is upregulated during drip-flow biofilm formation (20). We therefore sought to determine whether RsmW contributes to *P. aeruginosa* biofilm formation under iron limiting conditions. For this, we grew PAO1 and the isogenic Δ*rsmW* strains in flow-cell biofilms using 1% LB with or without 5 µM iron supplementation for 48 h. We observed similar biofilm formation of both PAO1 and Δ*rsmW* mutant when grown in iron-supplemented medium. However, under iron limiting conditions, Δ*rsmW* mutant exhibited a significant defect in biofilm formation compared to PAO1 (Fig. 6A-B and Fig. S4A). This phenotype was complemented *in trans* by the introduction of the *pa4570_rsmW* locus at the chromosomal CTX phage site (RsmW^C^) (Fig. 6C-D and Fig. S4B). These data demonstrate that RsmW plays an important role in biofilm growth under low iron conditions.

**Figure 6.**
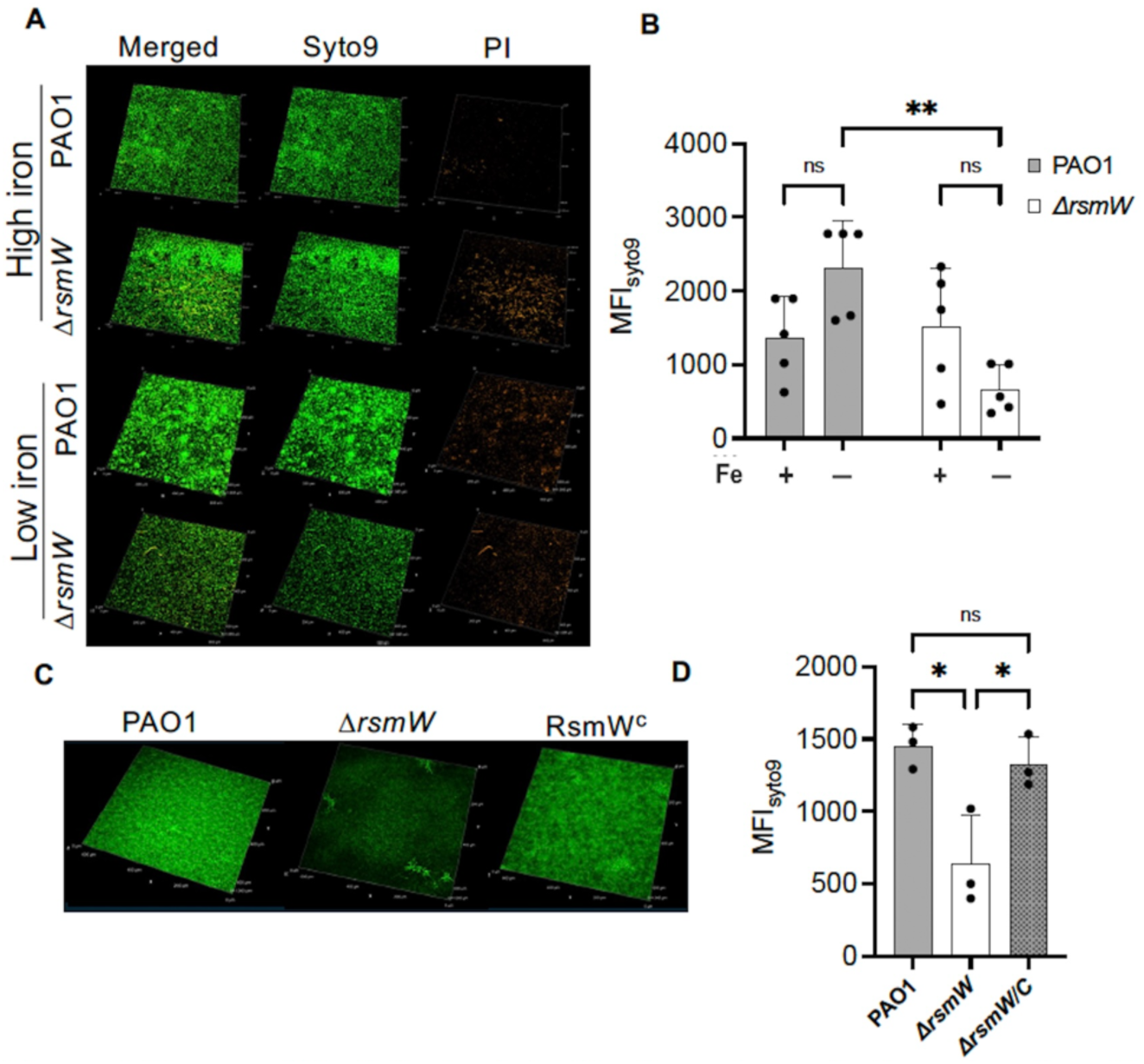
The Δ*rsmW* is defective in biofilm formation under iron starvation. Strains were grown in flow-cell biofilms with or without 5 µM iron supplementation as indicated. A-B) Representative merged confocal microscopy images (A) and quantitation of the max fluorescent intensity (MFI) (B) of wild type PAO1 and Δ*rsmW* (n= 5). C-D) Representative merged confocal images (C) and MFI quantitation (D) of PAO1, Δ*rsmW* and RsmW^C^ (n=3). SYTO9 (green) was used for staining live cells and propidium iodide (red) was used for staining eDNA. The red fluorescence of propidium iodide has been pseudo-colored with orange. MFI results are represented as mean ± SD; each biological replicate is represented a black circle data point. Asterisks indicate a statistical difference as determined by two-way (B) or one-way (D) ANOVA followed by Bonferroni’s multiple comparisons test: * P < 0.05, ** P < 0.005. Images of all biological replicates are shown in the Supplementary Materials (Fig. S4A-B).

### RsmW is required for full virulence in a *G. mellonella* infection model

Since the sequence for RsmW is found in *P. aeruginosa* but not in non-pathogenic *Pseudomonas* species, we sought to determine whether RsmW would impact *P. aeruginosa* pathogenesis in an animal model. To this end, we used *G. mellonella* larvae, an infection model that recapitulates murine infection phenotypes of many different *P. aeruginosa* mutant strains and, similar to the mammalian immune system, induces metal sequestration responses upon infection (32–35). *G. mellonella* larvae were infected with either 10 or 100 colony forming units (CFU) of PAO1 or the isogenic Δ*rsmW* and RsmW^C^ strains, and survival of the infected larvae was monitored for 18 h. We observed delayed larval killing when infected with 10 CFUs of the Δ*rsmW* mutant compared to PAO1 and RsmW^C^ (Fig. 7, and Fig. S5A-C). These data suggest that metal sequestration by *G. mellonella* leads to RsmW-dependent induction of virulence traits.

**Figure 7.**
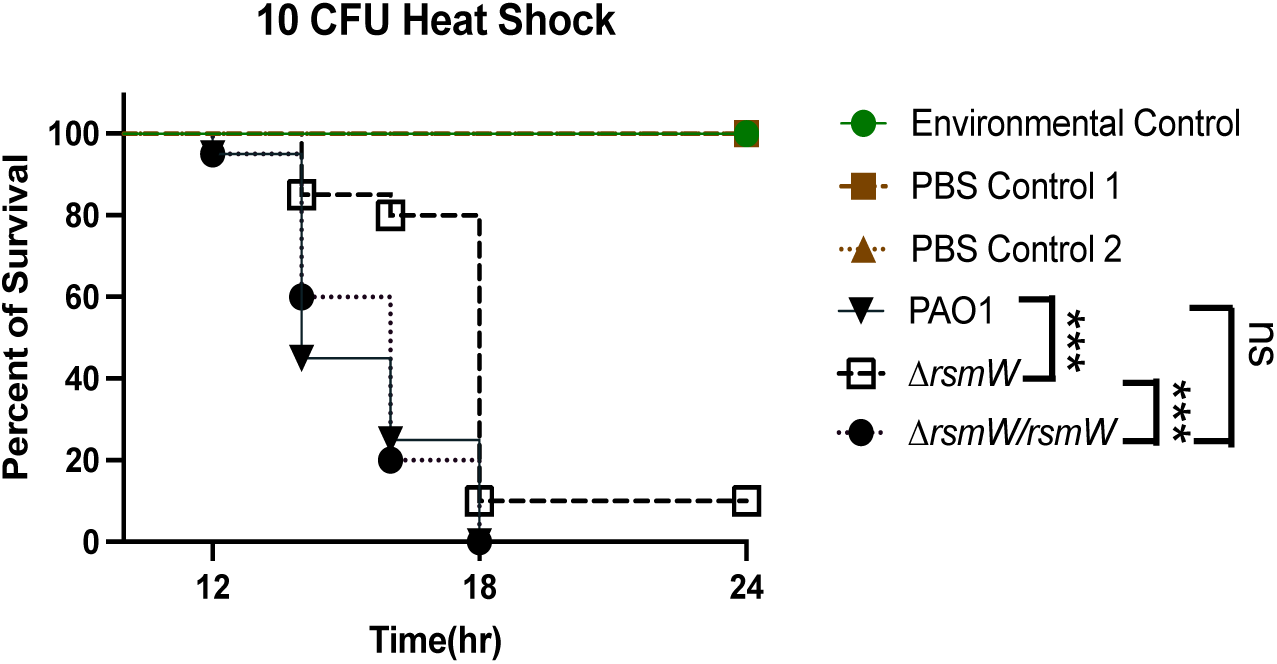
The Δ*rsmW* strain is attenuated *Galleria mellonella* larvae infection model. Representative survival curve of *G. mellonella* larvae infected with 10 CFU of the indicated strains. Strains were grown in LB to an OD_600_ of 0.5, diluted in PBS and injected into the larval hemolymph via the last proleg. 10 larvae were used for each of the controls and 20 larvae were used for infection by each strain. All the larvae were incubated at 37°C and their survival were monitored till 24 h. Results are represented as Kaplan-Meier plots representing larvae survival percentage over time following infection. Asterisks indicate statistical significance between indicated strains showing pairwise comparisons plotted by Bonferroni’s multiple comparisons test: *** P < 0.0005. Survival plots for each infection experiment are shown in the Supplementary Materials (Fig. S5A-C).

### Iron regulates *pa4570_rsmW* in lung infection isolates from individuals with CF

The Rsm sRNAs are associated with shifting from acute to chronic virulence traits, known to be hallmark of CF lung infection (26,31,36,37). We therefore determined whether iron regulates expression of *pa4570_rsmW* in *P. aeruginosa* strains isolated from sputa of CF patients. Three CF isolates––termed JSRI-1, JSRI-2, JSRI-3 to indicate their order of isolation from an individual person with CF––were selected due to their distinct colony morphologies despite being clonal to one another (18). All three isolates showed significant induction of RsmW sRNA levels in low iron conditions compared to high iron conditions (Fig. 8). Thus, iron regulates the expression of *pa4570_rsmW* in these clinical isolates despite their isolation from different stages of CF lung infection.

**Figure. 8.**
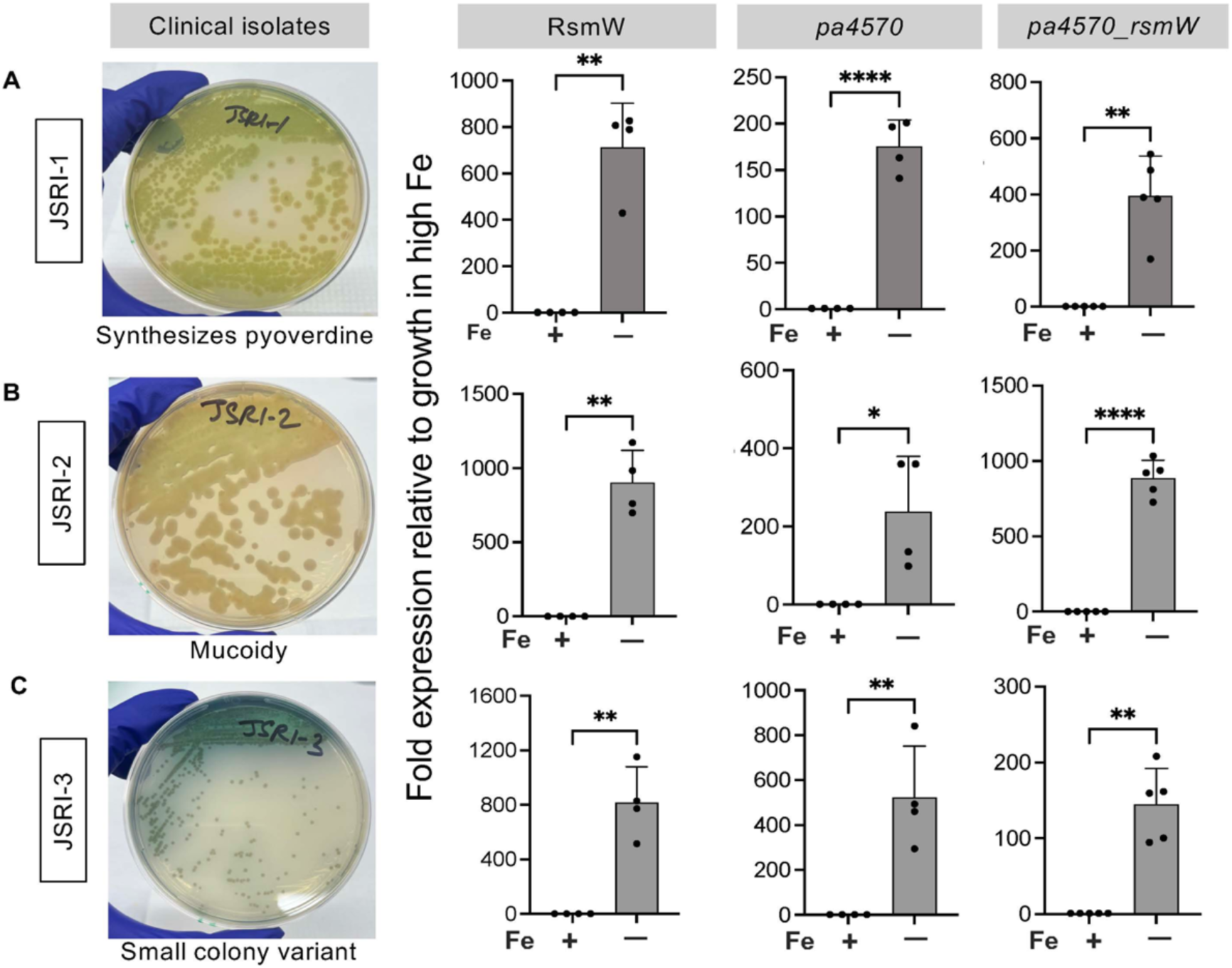
Iron regulates expression of *pa4570_rsmW* in clonal, longitudinal CF isolates that exhibit distinct phenotypes. CF isolates JSRI-1 (A), JSRI-2 (B), and JSRI-3 (C) were streaked onto LB agar plates to ensure their previously reported phenotypes (ref). Each isolate was then grown in DTSB media with or without iron supplementation for 18 h. Cultures were harvested, and RNA was isolated for cDNA synthesis and qPCR analysis. At least 4 biological replicates and 3 technical replicates per biological replicate were analyzed. Each biological replicate is represented a black circle data point. Results are represented as mean ± SD. Asterisks indicate a statistical difference as determined by T-test: * P < 0.01, ** P < 0.005, **** P < 0.0001.

## DISCUSSION

Iron likely plays a crucial role in the patho-adaptation of *P. aeruginosa* to the CF lung as chronic infection and inflammation lead to decreased pulmonary function. However, the mechanisms that underly this adaptation remain poorly understood. In this study, we report a novel link between iron and the Rsm regulatory system, which mediates the shift from acute to chronic virulence gene expression during CF lung infection. We demonstrate that iron starvation results in robust induction of RsmW, and we provide evidence for RNAse E mediated cleavage of the RsmW sRNA from the iron-regulated *pa4570* transcript. We further show that RsmW is required for biofilm formation in iron-depleted conditions as well as for full virulence in *G. mellonella* larvae. Lastly, we demonstrate that iron-dependent regulation of RsmW is maintained in CF isolates form various stages of lung infection. Combined, these findings present a novel and intriguing role for RsmW in iron dependent adaptation of *Pa* from acute to chronic infection phenotypes in the CF lung.

We previously showed that two well-characterized Rsm sRNAs, RsmY and RsmZ, are induced upon iron limitation (19). However, this regulation was modest and occurred only under static conditions in a PqsA- and PqsR-dependent manner (19). In contrast, iron regulation of RsmW likely occurs via a predicted Fur binding site upstream of *pa4570*. Fur also regulates the PrrF sRNAs that are central for maintaining iron homeostasis during low iron growth. In contrast to the PrrF sRNAs, loss of RsmW does not result in a growth defect when *P. aeruginosa* is grown in low iron media. This stark contrast in growth phenotypes suggests that RsmW contributes less to iron homeostasis and more as a regulator of virulence phenotypes. While *pa4570* is conserved across all the Pseudomonads, RsmW is specific to *P. aeruginosa* (Fig. S1), and it is induced at 37°C (20). This suggests that RsmW was acquired to convey iron- and temperature-dependent expression of virulence traits during mammalian infection. Although not a mammalian model, a recent study showed that deletion of *retS* significantly attenuated virulence of PAO1 in *G. mellonella* (38).This result agrees with our findings as *retS* deletion results in reduced RsmW expression (Fig. 8). Furthermore, our findings also showed that RsmW mediates *P. aeruginosa* biofilm formation, which is a major contributor to chronic virulence. Presumably these phenotypes occur via RsmA since, similar to other Rsm sRNAs, RsmW binds to and sequesters RsmA (20). Whether *pa4570* is also associated with biofilm formation and virulence remains to be determined, though the PA4570 has been detected previously under iron starvation conditions (16). The N-terminus of PA4570 is similar to that of the RNA binding Rsm proteins (20), highlighting the potential for this protein to participate in RsmW-dependent regulation of *P. aeruginosa* virulence (20). Elucidation of the mechanisms by which RsmW and PA4570 regulate virulence in *P. aeruginosa* warrants further investigation.

The pronounced difference in stability between *pa4570* and RsmW prompted us to hypothesize that RsmW is processed by RNAse E, an endoribonuclease that mediates RNA processing events (39,40). The Δ*rne 50bp c-term* mutant strain showed a significant decrease in the levels of RsmW, which correlated with increased levels of the *pa4570*_*rsmW* co-transcript. The same mutant also showed increased RNA levels of total *pa4570*. One plausible explanation for this finding is that the unprocessed transcript is more stable than processed *pa4570* due to protection of its 3’ end by the RsmW sequence, as RNAse E mediated cleavage commonly promotes mRNA decay in *Escherichia coli*, *Salmonella enterica*, and *P. aeruginosa* (41). However, RNA levels of *pa4570* are only slightly and not significantly reduced in the *rsmW* deletion mutant, arguing against this likelihood. Together, these data underline the need to better define the relationship between RNA processing and stability of *pa4570* and RsmW, as well as how RNAse E contributes to these processes. Previously, enteric pathogens have been shown to have 3’UTR derived sRNAs generated via the endonuclease activity of RNAse E followed by other processing events involving factors such as the Hfq chaperone and the exoribonuclease polynucleotide phosphorylase, PNPase (41). RNAse E and PNPase have also been reported to contribute to *P. aeruginosa* virulence (39,42,43).The detailed characterization of interactions between RsmW and RNAse E, and virulence regulation associated with these interactions, remain to be explored in future investigations.

Combining all our findings, we conclude that RsmW contributes to iron dependent regulation of chronic virulence phenotypes in *P. aeruginosa*. This study further demonstrates that upstream regulation of RsmW is distinct from other Rsm sRNAs and suggests that the impact of distinct Rsm sRNAs on virulence depends on both transcriptional and post-transcriptional regulatory events. This idea is supported by a prior report showing differential expression of the Rsm sRNAs at different growth stages of *P. aeruginosa* growth (24). Indeed, determining how the Rsm network integrates various host signals is critical to understanding how *P. aeruginosa* adapts to different infection sites. Taken together, this work establishes a novel line of investigation into how host-mediated iron sequestration drives *P. aeruginosa* pathogenesis.

## MATERIALS AND METHODS

### Bacterial strains and growth conditions

All the bacterial strains that are used in this study are listed the supplementary file (Table S1). *P. aeruginosa* lab strains (PAO1 and PA14) and the clonal isolates (JSRI-1, JSRI-2, JSRI-3) were routinely grown overnight by streaking from freezer stocks in tryptic soy agar (TSA) (Sigma, St Louis, MO) plates having 15 g/L agar. For experiments, five isolated colonies of each strain were inoculated in tryptic soy (TSB) broth from the streaked TSA plates. Cultures were grown in Chelex-treated and dialyzed trypticase soy broth DTSB medium supplemented with either 0 or 100 μM FeCl_3_ as previously mentioned (ref). Cultures were incubated in shaking conditions for 18 h at 37°C. The antibiotics were added in the following concentration: ampicillin, 100 μg/ml; tetracycline, 10 or 15 μg/ml, for *E. coli* and carbenicillin, 250 μg/ml and tetracycline, 150 μg/ml for PAO1 in case of genetic manipulations.

### Generation of deletion strains

Deletion mutants were generated according to previously published protocol with specific modifications (19). Briefly, genomic DNA were isolated and flanking sequences for the target gene were fused in pEXTc18 suicidal vector by Gibson method as mentioned previously (19). The *retS* deletion construct was generated in pMQ30 vector was provided by Prof. Katharina Ribbeck lab at MIT (29). The deletion constructs were then introduced into the PAO1 genome followed by replacement of the native loci of the target gene via allelic exchange. Unmarked deletions were screened by PCR using primers mentioned in the supplementary file (Table S1).

### RNA Isolation

Bacterial cultures were pelleted by centrifugation and stored in RNA latter at −80°C for long-term. RNA isolation was performed as previously described (19). In short, bacterial cells were resuspended in TE buffer with lysozyme followed by lysis with RLT-buffer of the RNeasy kit and column based purification method. An additional DNase treatment (NEB DNAse) was performed when required. Purified RNA was subjected to overnight precipitation with Sodium acetate and 100% ethanol. Finally, pure RNA was resuspended in nuclease-free water.

### Real time PCR and northern blot analysis

50 ng of RNA was used for cDNA synthesis as mentioned earlier (13). The cDNA was diluted 1:10 times and used for real time PCR. Northern blot was performed as previously described (13). Briefly, RNA was run in 10 % urea-polyacrylamide gel followed by transfer to nylon membrane and treated with custom made biotinylated probes for RNA detection. All the primers and probes are mentioned in the supplementary table (Table S2).

### Biofilm growth

Biofilms were grown as previously mentioned (44). Briefly, bacterial culture was inoculated in flow cells. These flow cells were connected to a peristaltic pump that maintains a constant flow of media that are connected to the flow cells, incubated for 48 h at 37°C.The media used is 1% LB and metals (Cu, Zn, Mn and Ni) but with or without supplementation of 5 µM Fe. Biofilms were stained with Syto9 for live cells and propidium iodide (PI) for dead cells and eDNA. The biofilm growth was imaged using Laser scanning confocal microscopy and analyzed by ImageJ.

### Galleria mellonella survival assay

*G. mellonella* larvae were obtained from a local vendor. The last instar larvae stage was chosen, and survival assay was performed with slight modification of the method as described by other groups (35). *P. aeruginosa* cultures were diluted to 1:10 in LB and subcultured to an OD_600_ of 0.5. Bacteria were then serially diluted to 10^4^ and 10^3^ CFU in PBS. For each strain, groups of 20 larvae, weighing between 200-300 mg were used. All the larvae were heat shocked at 37°C prior to infection with *P. aeruginosa* cultures. This has been shown to increase immunity in the larvae (45). 10 µl of bacterial suspension was injected into each of the larval hemolymph through the last right proleg using a Hamilton micro-syringe equipped with a 30-gauge needle (Hamilton Company). A group of 10 larvae were chosen for injecting PBS as an infection control. Two other groups were also included for environmental control and heat-shock control.

### Statistical analysis

Statistical analyses were performed in all the experiments by using two-way ANOVA on Prism 9, using Bonferroni’s post-test for multiple comparisons, with a significance threshold of *p* value < 0.05. Real time experiments were conducted as previously mentioned (19). Briefly, each experimental data has at least three biological replicates and three technical replicates the results were represented as mean ± SD. All the data points represent biological replicates. In case of the *G. mellonella* infection model, data analysis was performed by Kaplan Mayer plots and results were represented as mean ± SD. Significance was calculated by two-way ANOVA with post-tests for multiple comparisons.

## ACKNOWLEDGEMENT

We are grateful to Drs. Susana Mouriño and Angela Walks for generously sharing the PAO1/Δ*rne 50bp c-term* mutant. This work was funded by NIH grants R01AI123320 and R01AI161294 (to AGO) and the University of Maryland School of Pharmacy.

## Notes

### Competing Interest Statement

The authors have declared no competing interest.

